# Learning when to learn: hummingbirds adjust their exploration behaviour to match the value of information

**DOI:** 10.1101/2023.09.08.556950

**Authors:** C. Donkersteeg, H. Peters, C. Jerome, C. Menzies, K. C. Welch, L. M. Guillette, R. Dakin

## Abstract

Exploration is a key part of an animal’s ability to learn. The exploration-exploitation dilemma predicts that individuals should adjust their exploration behaviour according to changes in the value of information. Here, we test this prediction by tracking ruby-throated hummingbirds as they foraged repeatedly from a large array of artificial flowers, wherein 25% of the flowers contained a sucrose reward. Similar to real-world floral dynamics, the reward locations were consistent in the short-term, but varied from day to day. Thus, the value of information about the flower contents would be greatest at the beginning of a daily foraging session, and decay toward the end of each session. We tracked five individual hummingbirds in repeated foraging sessions, comprising more than 3,400 floral probes. We analyzed two metrics of their exploration behaviour: (1) the probability that a bird would shift from probing one flower to another, and (2) the Shannon information entropy of a sequence of flowers probed. We show that initially, the hummingbirds increased their exploration behaviour as time elapsed within a session. As they performed more sessions and learned the rules of the environment, the hummingbirds switched to explore more diverse choices at the beginning of a foraging session, when the value of information was high, and less diverse choices toward the end of a session. Our results suggest that foraging hummingbirds can learn when to learn, highlighting the importance of plasticity in exploration behaviour.

**Highlights:**

- Exploration is a necessary part of learning
- Foragers must balance sampling for information with the use of known rewards
- Hummingbirds learned to explore more when the value of new information was high
- Apparent mistakes may actually represent an information-seeking strategy

## 1. Introduction

Exploration is a key part of learning, and is broadly defined as behaviour that gathers information (e.g., Reader 2015). In nature, individuals often face a dilemma between exploring a new stimulus or exploiting a familiar one (Gopnik, 2020; Krebs et al., 1978). Theoretical modelling of this exploration-exploitation dilemma has led to advances across diverse fields, including ecology, psychology, neuroscience, and computing (Gopnik, 2020; Kaelbling et al., 1996; O’Farrell et al., 2019; Schulz & Gershman, 2019). A key assumption of exploration-exploitation models is that individuals are expected to adjust their exploration behaviour according to the costs and benefits of information-seeking, exploring more when the value of new information is high. In animals, this tuning of exploratory behaviour can occur as a result of plasticity (i.e., by learning when to explore) or heritable genetic change (Sherratt & Morand-Ferron, 2018).

Despite its fundamental theoretical importance, exploration is often overlooked in empirical studies of animal cognition and behaviour. In order to determine what animals know or what they can detect, researchers in cognition and behaviour often use paradigms that attempt to extinguish exploratory tendencies (e.g., via food deprivation) (Shettleworth, 2010). Under these paradigms, repeated visits to non-rewards are often assumed to be mistakes in recall (Bateson et al., 2003; Healy & Hurly, 1995; Vámos et al., 2020); however, this behaviour may also represent an exploratory foraging strategy. In nature, many resources are always dynamic in space and time. For example, flowers produce nectar at different concentrations, rates, times, and locations, and the flowers producing nectar also change as seasons progress (Berger-Tal & Bar-David, 2015; Henderson, Hurly, Bateson, et al., 2006; Tello-Ramos et al., 2019). Nectar within a single flower can also replenish within hours or minutes, requiring regular updates to keep track of profitability. In these dynamic environments, individuals are expected to accumulate more resources in the long-term if they can adjust their exploration behaviour based on the value of information in their environment (O’Farrell et al., 2019).

Nectar-feeding hummingbirds are a useful group for studying exploration, because they require frequent nectar meals multiple times per hour to fuel their high metabolic rates (López-Calleja et al., 1997; Suarez, 1992). Given the small amounts provided by each flower, hummingbirds often need to track the locations of numerous flowers, sometimes kilometres apart, to meet their energetic needs (Gass et al. 1976). Hummingbirds have evolved sophisticated spatial memory and learning ability, such that they can quickly acquire information about a landscape of floral rewards (González-Gómez et al., 2011; Tello-Ramos et al., 2015, 2019; Wolf & Hainsworth, 1990). They are able to track both the location and timing of their nectar food sources (Ayers et al., 2018; Bateson et al., 2003; Biernaskie et al., 2002; Henderson, Hurly, Bateson, et al., 2006; Heyneman, 1983; Morgan et al., 2014; Temeles et al., 2006). The hippocampal formation is also greatly enlarged in hummingbirds, in support of their reliance on enhanced spatial memory (Ward et al., 2012).

Here, we examine exploration behaviour of ruby-throated hummingbirds, and test if the birds adjust their exploration behaviour based on changes in the value of information in a dynamic foraging environment. To do this, we tracked five captive individual hummingbirds as they foraged repeatedly from a large artificial flower array (Fig. 1A). Similar to real flowers, only a subset of the artificial flowers contained artificial nectar. The resource dynamics in our experiment were determined by a simple rule: reward locations were consistent in the short-term (within a day) but varied randomly among daily foraging sessions. This schedule meant that the value of information about a flower’s status would decay toward the end of each foraging session. Exploration-exploitation theory predicts that foragers that have learned these dynamics should explore more at the beginning of a foraging session and become less explorative toward the end of each foraging session. To test this prediction, we tracked each hummingbird across seven daily foraging sessions, providing each bird with the opportunity to learn how the floral dynamics worked. We then analyzed measures of the diversity and information content of the flowers they probed to test how the birds varied their exploration behaviour in response to this environment.

**Figure 1.**
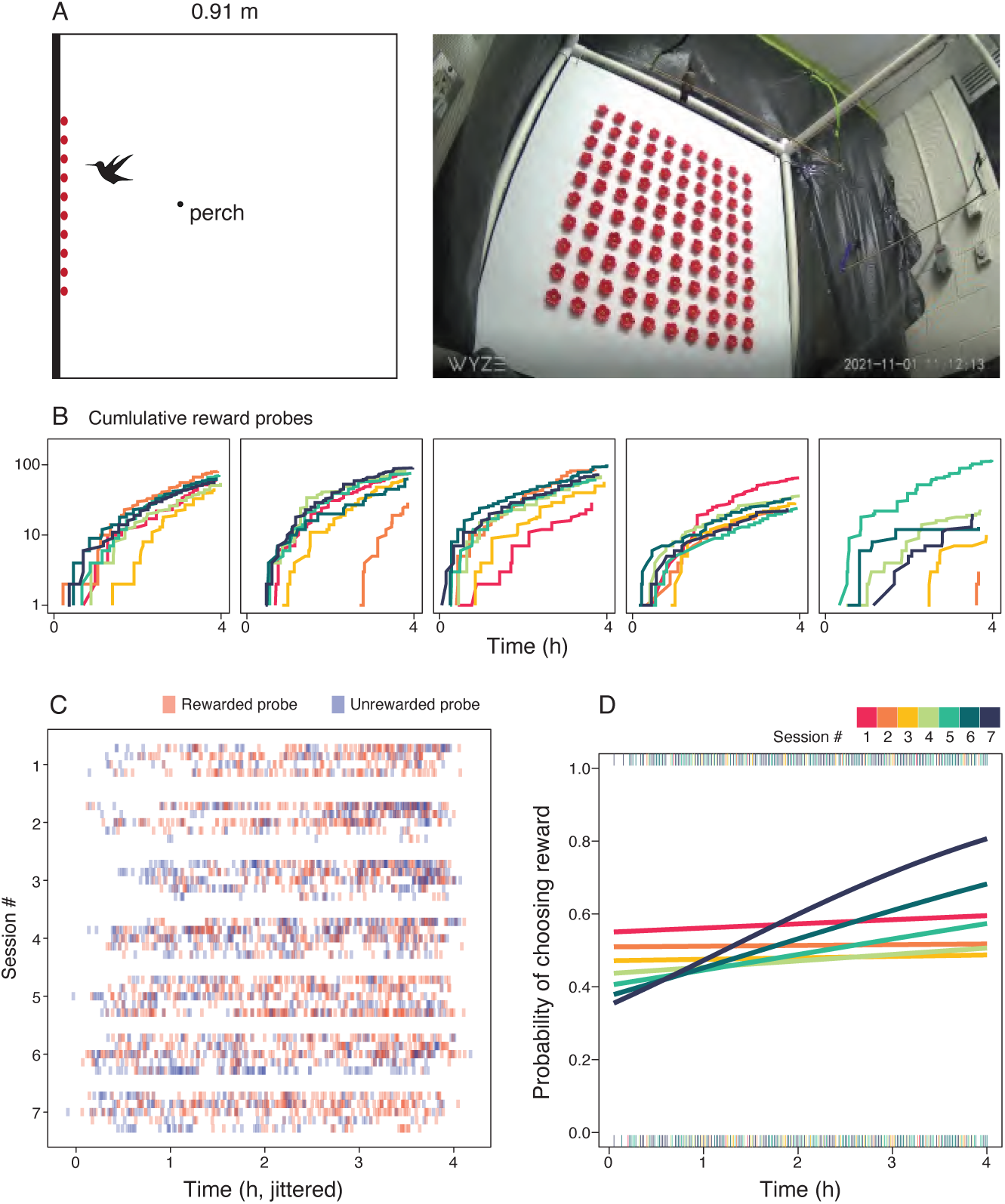
Hummingbirds sampled from a complex array of flowers. (A) Birds foraged from a 10 x 10 array of flowers in a cubic chamber over repeated sessions. (B) Cumulative probes to rewarded flowers over time. Each panel in (B) displays one bird; note that probe counts are represented on a log scale. The hummingbirds typically located their first rewarded flower within the first four probes of a session. Yet even after locating rewarded flowers, the hummingbirds continued to probe non-rewarded flowers frequently. The plot in (C) shows probes to rewarded (pink) and nonrewarded flowers (blue) through time. Each bird-session is shown in a row, with probes represented by semi-transparent bars. Probes are jittered slightly along the x-axis to visualize the general trends. (D) With increasing experience (i.e., by session 7), the hummingbirds were far more likely to probe rewarded flowers near the end of the session. Surprisingly, they were also less likely to probe rewarded flowers during the first hour of their latter trials. The solid lines in (B) show predictions from the fitted model of foraging performance.

## 2. Methods

### Study animals

We tested five wild-caught ruby-throated hummingbirds (three males and two females) that were captured on the campus of the University of Toronto Scarborough (43.783323, -79.189811). The birds were captured as adults at outdoor hummingbird feeders and were housed in captivity for at least two months prior to this study. These hummingbirds thus had extensive experience with real flowers in the wild (which vary in nectar content), as well as outdoor hummingbird feeders and feeders in captivity that provided food *ad libitum*. In the vivarium, each hummingbird was housed individually (0.9 x 0.5 x 0.5 m cages) in a room where other individuals hummingbirds could be seen and heard. Ambient conditions in the vivarium were held between 20-24 C and 30-70% humidity, with 13 hours light/11 hours dark to approximate light cycles where the birds would naturally occur.

### Ethical note

All procedures were approved by Carleton University (AUP #111381) and the University of Toronto Scarborough (AUP #20011649).

### Artificial flower array

The artificial flower array consisted of 100 flowers in a 10 x 10 square grid mounted on an opaque white board (Figure 1A). The board was mounted in a cube-shaped chamber (0.91 m^3^) of fine mesh netting, with a perch positioned 32 cm from the flower array at its midpoint elevation (Figure 1A). Each artificial flower had red plastic petals with a painted yellow center, designed to be consistent in appearance, with a central opening that provided access to a 1.5 mL pipette tube located behind the white board. Flower tubes were either filled with 0.5 mL of 20% sucrose solution (rewarded) or empty. We based this setup on the typical metabolic requirements for a ruby-throated hummingbird within a small chamber, where most of their time is spent perching. In preliminary tests of the array, we also confirmed that 0.5 mL per flower was a sufficient volume that flowers with artificial nectar would not be depleted over a 4-hour foraging session, while also encouraging frequent probing of the artificial flowers (Varma et al., 2020).

### Foraging sessions

Foraging sessions were conducted in October and November of 2021, which corresponds to the end of post-breeding migration and beginning of the overwintering period in this species. Individual foraging sessions took place in a dedicated room that was isolated from other birds and from the researchers. Each bird was placed in the chamber for seven 4-hour foraging sessions, with two to three days between each session. Captive hummingbirds typically feed at 5- to-10-minute intervals (Fig 1B; see also Dakin et al., 2016). In each session, 25% of the flowers in the array contained a food reward, with the reward configuration determined randomly for each session. The same random configuration was used for each of the five birds in their n^th^ session. Hence, the reward locations in the array were consistent in the short-term (within days) but varied across days (and across a given bird’s daily foraging sessions).

All artificial flowers were made to be functionally identical with no visual clues as to which ones contained rewards (and which were empty), other than their location. The pipette tubes that held the sucrose solution were not visible from within the chamber. Hummingbirds do not use odour to locate sucrose rewards (Goldsmith & Goldsmith, 1982; Núñez et al., 2021). Therefore, the birds in our experiment had to rely on exploration and short-term spatial memory to collect food. The spatial cues that the hummingbirds may have used included the position of flowers relative to the edges of the array, the features of the chamber and perch, and/or the features of the surrounding room (Fig 1A), as well as the ordinal position of flowers within the array (Vámos et al., 2020).

Before each foraging session, the bird’s body mass was recorded. During the session, behaviour was recorded using a motion-activated camera (Wyze Cam v3). We define a probe event as any insertion of the hummingbird’s bill into an artificial flower. We determined which flowers were probed, and at what times, from the video recordings. The hummingbirds had a strong bias toward using the upper rows of the array that were located above the height of the perch; for example, 93% of probes were made to the 30 flowers located in the top three rows of the array. This bias was not anticipated when randomly assigning the reward locations; on average, the proportion of rewarded flowers within the top three rows was 29% (averaged across the seven configurations). In other studies of related hummingbird species, the birds consistently maintain a position within the top 20-30 cm of their flight chambers and rarely descend near the bottom (Dakin et al., 2016), possibly because positions near the ground put hummingbirds at increased vulnerability to threats.

### Behavioural scoring

Our aim in this study was to measure exploration by each hummingbird how it changed with time and experience in the foraging sessions. To this end, we analyzed two metrics of exploration behaviour that capture the potential to acquire new information without making any assumptions about an individual’s intentions or memory state (Shettleworth, 2010). First, we analyzed the probability that a bird shifted its choice of flower from one probe to the next. We consider shifting to be explorative, because shifting increases the potential information about reward distributions in time. As an example, the sequence ABABAB probes two flowers, A and B, with five shifts between them, whereas the sequence AAABBB probes the same two flowers, but has only one shift. All else being equal, when an individual shifts more frequently among different flowers, it is able to gain more recent information about the contents of different flowers. We analyzed the probability of shifting per each probe event, with further details provided in the section below.

As our second metric of exploration, we analyzed floral choice diversity integrated over time. To quantify this measure, we used the Shannon Index, a measure of entropy or information content in a distribution (Magurran, 2004; Shannon, 1948). Importantly, the Shannon Index accounts for both the number of unique members in a set and how evenly those items are represented in the set; this particular metric is also commonly used to estimate the diversity of ecological communities. In our case with flower choices, the two sequences ABBBBB and AAABBB both probe two unique flowers, but the sequence AAABBB will have a higher information content (greater entropy) because of its greater evenness. We consider foraging sequences with greater Shannon entropy to be more explorative because they increase the amount information gained for all locations. We used the vegan 2.6-2 package (Oksanen et al., 2022) to compute the Shannon Index (SI) for a bird’s most recent set of five probes (up to and including a given probe). We chose to integrate over 5 probes because this provided a balance between precision in the SI estimates and our ability to resolve how these estimates changed over time within sessions. Note that SI computed for very short sequences is poorly estimated; SI does not exist for a set of one. As a check on analyses involving choice diversity, we also verified that all conclusions were unchanged if SI was computed using alternative sequence lengths between 4-10 probes.

### Data analysis

All data processing and analyses were performed in R 4.2.2 (version 4.2.2., R Core Development Team 2020). We analyzed three aspects of foraging behaviour: (1) the probability that a bird chose a rewarded flower, on a per probe basis (binary yes/no); (2) the probability that a bird shifted its choice from the previous flower, on a per probe basis (binary yes/no), and (3) the choice diversity of a bird’s most recent five probes, on a per probe basis. The first analysis (1) can be considered an analysis of foraging performance, whereas (2) and (3) represent our two measures of exploration behaviour in the flower array.

For each of the these three response variables, we fit a mixed-effects regression model using the packages lme4 1.1-31 (Bates et al., 2023) and lmerTest 3.1-3 (Kuznetsova et al., 2020). The fixed effect predictors included session number (1-7) as a measure of an individual’s experience of the experimental paradigm, the time elapsed since the start of the session, and the interaction between session number (experience) and time elapsed. This interaction allowed us to test whether foraging behaviour within sessions depended on a bird’s accumulated experience of the environment. We modelled session number as a 2^nd^-order polynomial to allow for nonlinear change with increased experience. All models also included a fixed effect of body mass, as well as a random effect of bird ID to account for repeated measures.

The two binary response variables (the probability of choosing a reward, and the probability of shifting between flowers) were modelled with a binomial error distribution. Choice diversity was modelled with a Gaussian distribution. We omitted one individual’s first session from these analyses because that bird did not probe the array for >2 hours during its first session. We consequently recorded that bird with the session 1 reward configuration again the following day. Because that bird’s session 1 was not performed contiguously, we removed it from further analyses. The sample size for the analysis of reward probability was 3,433 probes across 34 foraging sessions by five birds (average of 101 probes per session ± 43 SD). Of these, 3,399 probes represent the sample size for the analysis of choice shift (because shift is quantified for probes occurring after the 1^st^ probe in a given session), and 3,297 probes represent the sample size for the analysis of choice diversity (because diversity is quantified for probes occurring after the 4^th^ probe in a given session).

## 3. Results

### (a) Description of flower use in the array

The hummingbirds frequently probed non-rewarded flowers in the array, even after they had repeatedly located multiple rewarded flowers. By the 2-hour midpoint of the sessions, the hummingbirds had on average discovered 4.7 unique rewarded flowers (± 1.9 SD) and accumulated 20.7 reward probes (± 11.2 SD; Figure 1B). From that midpoint time until the end of the sessions, the birds continued to visit unrewarded flowers 56% of the time, on average (Figure 1C).

We checked whether the birds initially chose flowers that they had experienced as rewarded in their previous session (see supplement for details). The birds usually (62% of the time) began the sessions by probing flowers that were not experienced as rewards in their last session. They were slightly more likely to initially probe previous rewards than expected by chance (38% actual probability vs. 20% chance expectation; paired t-test p < 0.0001, n = 30 bird-sessions).

### (b) Probability of choosing a rewarded flower

When analyzing the probability of choosing a reward, there was a statistically significant session-by-time interaction (LRT p < 0.0001; see Figure 1D and Table S1). In their first session, the birds were relatively stable in the proportion of probes made to rewarded (vs. unrewarded) flowers (see the line for session 1 in Figure 1D). By their 7^th^ session, the hummingbirds were far more likely to probe rewarded flowers by the end of the session (session 7 in Figure 1D). Surprisingly, they became significantly *less* likely to probe rewarded flowers during the first hour of the sessions (e.g., session 7 estimate of 36% probability [95% CI 28-44%] for the first hour, vs. session 1 estimate of 54% probability [95% CI 45-67%] for the first hour).

### (c) Probability of shifting between flowers

When analyzing the probability of shifting between flowers, there was a significant session-by-time interaction (LRT p < 0.0001; see Figure 2A and Table S1). In the initial sessions, the hummingbirds were unlikely to shift flowers early in the session, but then they became more likely to shift flowers as time elapsed (see the line for session 1 in Figure 2A). As birds gained experience with the array, this pattern reversed: they became more likely to shift between flowers at the beginning of a session, and less likely to shift between flowers at the end of a session (session 7 in Figure 2A).

**Figure 2.**
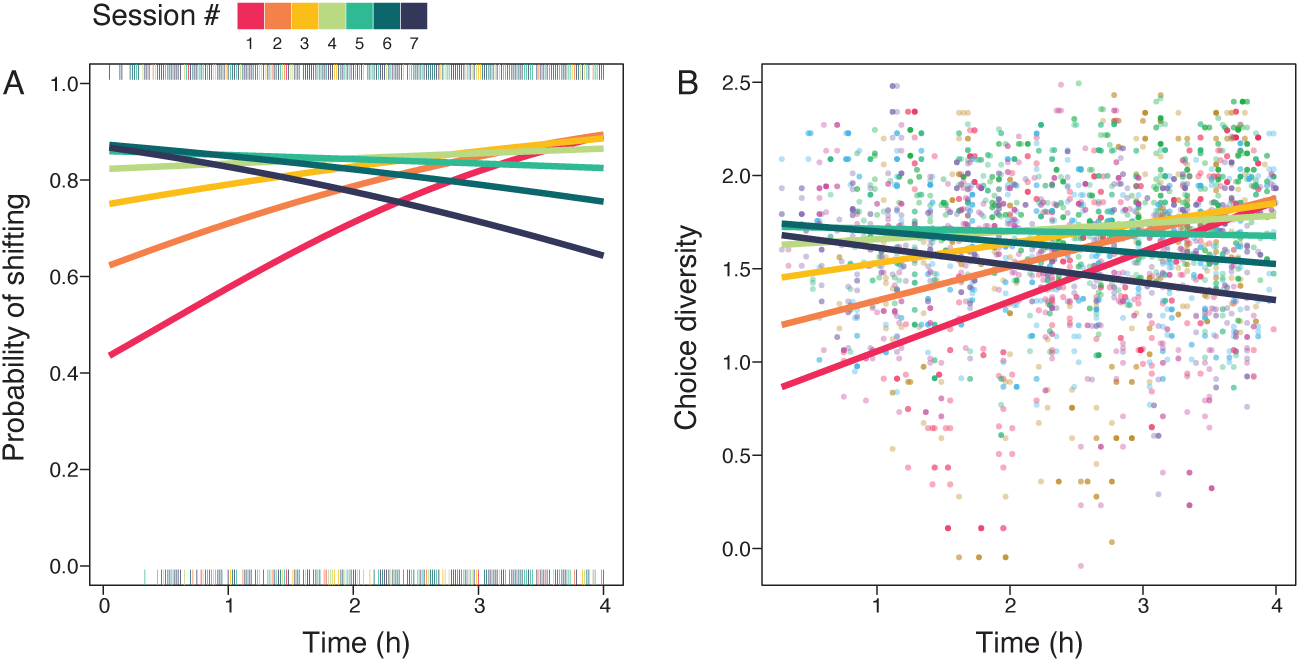
Hummingbirds learned when to explore. We analyzed two metrics of exploration: (A) shifting between flowers from one probe to the next, and (B) the diversity of recent flowers probed. The solid lines show predictions from fitted models. At the beginning of the experiment (i.e., session 1), hummingbirds scored low on both measures of exploration, but they became more explorative as their initial session continued. By the end of the experiment (session 7), they explored much more at the beginning of a session, and much less toward the end.

### (d) Floral choice diversity

When analyzing floral choice diversity, we observed the same significant session-by-time interaction described for shifting behaviour (LRT p < 0.0001; see Figure 2B and Table S2). In their initial sessions, the birds became less explorative as time elapsed within the session. By the end of the experiment, the birds began to make more diverse choices at the beginning of a session, and less diverse choices by the end of the session.

## 4. Discussion

In many studies of animal cognition, researchers often omit or seek to extinguish behaviour that fails to exploit a reward. These nonrewarded choices are often assumed to be mistakes. Yet birds are known to sophisticated sampling and exploration behaviours when foraging for food (Krebs et al., 1978). Here, we studied hummingbird sampling under conditions where food was always available *ad libitum*, and we did not attempt to motivate birds to perform only exploitation, nor did we attempt to extinguish exploration. Under these conditions, the hummingbirds frequently probed non-rewarded flowers even after they had repeatedly located rewards. In our experimental environment, the value of information about reward locations would decay toward the end of each session. We found that, with increased experience of these dynamics, the hummingbirds began to sample more diverse choices earlier in the session when the specific reward locations were unknown, and less diverse choices toward the end of each session (Figure 2). These findings indicate that individual birds can plastically adjust their exploration behaviour to match changes in the value of information (Reader, 2015; Sherratt & Morand-Ferron, 2018). Our results also suggest that we should not assume that nonrewarded choices are random or errors; instead, they may often represent strategic decisions that are shaped by an individual’s priors for a given environment.

Based on these results, we suggest that hummingbirds can learn when to learn. Is this ability specific to hummingbirds? As nectarivores with high metabolic rates, hummingbirds rely on spatiotemporal memory and learning to forage successfully in the wild. Wild hummingbirds often rely of numerous flower patches that can be hundreds of metres or even kilometres apart, each of which may have its own nectar dynamics (Gass et al. 1976). Field and laboratory studies show that hummingbirds can readily learn the spatial location and quality of multiple nectar rewards (Bateson et al., 2003; Healy & Hurly, 1995; Henderson, Hurly, Bateson, et al., 2006; Henderson, Hurly, & Healy, 2006). Moreover, Henderson et al. (2006) showed that hummingbirds avoid revisiting recently sampled flowers and exhibit fundamental aspects of episodic memory in their routine foraging behaviour. This previous work demonstrate that hummingbirds have evolved sophisticated spatiotemporal memory as a result of their reliance on widespread and ephemeral food sources. We expect that this foraging niche and reliance on highly variable nectar sources would also favour the development of exploration plasticity.

An additional question is whether “learning when to learn” requires awareness of one’s own knowledge state. Metamemory, defined as the ability to judge one’s own learning, has primarily been considered in only a few nonmammalian species (Basile et al., 2015; Goto & Watanabe, 2012; Inman & Shettleworth, 1999; Jozefowiez et al., 2009; Shea & Heyes, 2010; Shettleworth, 2010). In birds, some corvids have demonstrated a capacity to monitor the strength of their memory before making choices in a controlled task (Goto & Watanabe, 2012). It is important to note that our experiment does not test for metamemory, and the hummingbirds could have adapted to these environmental dynamics without necessarily being aware of their own knowledge of any particular flowers. For example, hummingbirds may have a generalized response to perform more exploration whenever they are in new environments. The birds in our experiment may have learned that the experimental environment was “new” at the beginning of each session.

Hummingbirds, as well as many other animals, can internally track time intervals (Potts et al. 2004; Henderson et al. 2006; Shettleworth 2009; Tello-Ramos et al. 2019). Wild hummingbirds can also use both time of day and the order of events to inform their foraging decisions (Tello-Ramos et al. 2015). In our experiment, the behavioural shifts that we observed within foraging sessions may indicate the use of an internal clock once the birds had learned the 4-hour duration of the sessions. Future experiments could expand on our results here by testing whether hummingbirds alter their exploration-exploitation dynamics during varied-length foraging sessions, by training birds to associate the onset of a specific cue (such as a coloured light) with a time horizon (such as 30 minutes until the end of the session). Based on our results here, we predict that hummingbirds would learn to reduce their exploration, and increase exploitation, upon the onset of time horizon cues.

Most studies of animal learning have focused on individual differences in the tendency to explore quickly; in some bird species, these between-individual differences in explorative behaviour are also associated with individual differences in flexibility in learning tasks (Guillette et al., 2011; O’Hara et al., 2017). Our results here show that individual hummingbirds can adjust how they sample flowers, gathering more information when the value of new information is highest. This same theoretical principle may also explain why animals (including humans) become less explorative and less willing to interact with novel stimuli as they age, as well as other forms of plasticity in exploratory behaviours (Reader, 2015; Sherratt & Morand-Ferron, 2018). Overall, our results highlight the importance of within-individual plasticity in information-seeking. By omitting or extinguishing exploration behaviour, we may be missing a major source of cognitive adaptation and diversity.

## Acknowledgements

We thank Ed Donkersteeg, Giulia Rossi, and the University of Toronto Scarborough Animal Care staff for help setting up the experiments, and Sue Bertram and Tom Sherratt for their valuable insights on this study.

## References

Ayers, C. A., Armsworth, P. R., & Brosi, B. J. (2018). Statistically testing the role of individual learning and decision-making in trapline foraging. Behavioral Ecology, 29(4), 885–893. 10.1093/beheco/ary058

Basile, B. M., Schroeder, G. R., Brown, E. K., Templer, V. L., & Hampton, R. R. (2015). Evaluation of seven hypotheses for metamemory performance in rhesus monkeys. Journal of Experimental Psychology: General, 144(1), 85–102. 10.1037/xge0000031

Bates, D., Maechler, M., Bolker [aut, B., cre, Walker, S., Christensen, R. H. B., Singmann, H., Dai, B., Scheipl, F., Grothendieck, G., Green, P., Fox, J., Bauer, A., & simulate.formula), P. N. K. (shared copyright on. (2023). lme4: Linear mixed-effects models using “Eigen” and S4 (1.1-34) [Computer software]. https://cran.r-project.org/web/packages/lme4/index.html

Bateson, M., Healy, S. D., & Hurly, T. A. (2003). Context–dependent foraging decisions in rufous hummingbirds. Proceedings of the Royal Society of London. Series B: Biological Sciences, 270(1521), 1271–1276. 10.1098/rspb.2003.2365

Berger-Tal, O., & Bar-David, S. (2015). Recursive movement patterns: Review and synthesis across species. Ecosphere, 6(9), art149. 10.1890/ES15-00106.1

Biernaskie, J. M., Cartar, R. V., & Hurly, T. A. (2002). Risk-averse inflorescence departure in hummingbirds and bumble bees: Could plants benefit from variable nectar volumes? Oikos, 98(1), 98–104. 10.1034/j.1600-0706.2002.980110.x

Dakin, R., Fellows, T. K., & Altshuler, D. L. (2016) Visual guidance of forward flight in hummingbirds reveals control based on image features instead of pattern velocity. Proceedings of the National Academy of Sciences, 113(32), 8849–8854. 10.1073/pnas.1603221113

Gass, C. L., Angehr, G., & Centa, J. (1976) Regulation of food supply by feeding territoriality in the rufous hummingbird. Canadian Journal of Zoology, 54(12), 2046–2054. 10.1139/z76-238

Goldsmith, K. M., & Goldsmith, T. H. (1982). Sense of smell in the black-chinned hummingbird. The Condor, 84(2), 237–238. 10.2307/1367678

González-Gómez, P. L., Vásquez, R. A., & Bozinovic, F. (2011). Flexibility of foraging behavior in hummingbirds: The role of energy constraints and cognitive abilities. The Auk, 128(1), 36–42. 10.1525/auk.2011.10024

Gopnik, A. (2020). Childhood as a solution to explore–exploit tensions. Philosophical Transactions of the Royal Society B: Biological Sciences, 375(1803), 20190502. 10.1098/rstb.2019.0502

Goto, K., & Watanabe, S. (2012). Large-billed crows (Corvus macrorhynchos) have retrospective but not prospective metamemory. Animal Cognition, 15(1), 27–35. 10.1007/s10071-011-0428-z

Guillette, L. M., Reddon, A. R., Hoeschele, M., & Sturdy, C. B. (2011). Sometimes slower is better: Slow-exploring birds are more sensitive to changes in a vocal discrimination task. Proceedings of the Royal Society B: Biological Sciences, 278(1706), 767–773. 10.1098/rspb.2010.1669

Healy, S. D., & Hurly, T. A. (1995). Spatial memory in rufous hummingbirds *Selasphorus rufus*: A field test. Animal Learning & Behavior, 23(1), 63–68. 10.3758/BF03198016

Henderson, J., Hurly, T. A., Bateson, M., & Healy, S. D. (2006). Timing in free-living rufous hummingbirds, *Selasphorus rufus*. Current Biology, 16(5), 512–515. 10.1016/j.cub.2006.01.054

Henderson, J., Hurly, T. A., & Healy, S. D. (2006). Spatial relational learning in rufous hummingbirds (*Selasphorus rufus*). Animal Cognition, 9(3), 201–205. 10.1007/s10071-006-0021-z

Heyneman, A. J. (1983). Optimal sugar concentrations of floral nectars—Dependence on sugar intake efficiency and foraging costs. Oecologia, 60(2), 198–213. 10.1007/BF00379522

Inman, A., & Shettleworth, S. J. (1999). Detecting metamemory in nonverbal subjects: A test with pigeons. Journal of Experimental Psychology: Animal Behavior Processes, 25(3), 389–395. 10.1037/0097-7403.25.3.389

Jozefowiez, J., Staddon, J. E. R., & Cerutti, D. T. (2009). Metacognition in animals: How do we know that they know? Comparative Cognition & Behavior Reviews, 4. 10.3819/ccbr.2009.40003

Kaelbling, L. P., Littman, M. L., & Moore, A. W. (1996). Reinforcement learning: A survey. Journal of Artificial Intelligence Research, 4, 237–285. 10.1613/jair.301

Krebs, J. R., Kacelnik, A., & Taylor, P. (1978). Test of optimal sampling by foraging great tits. Nature, 275(5675), 27–31. 10.1038/275027a0

Kuznetsova, A., Brockhoff, P. B., Christensen, R. H. B., & Jensen, S. P. (2020). lmerTest: Tests in linear mixed effects models (3.1-3) [Computer software]. https://cran.r-project.org/web/packages/lmerTest/index.html

López-Calleja, M. V., Bozinovic, F., & Martinez del Rio, C. (1997). Effects of sugar concentration on hummingbird feeding and energy use. Comparative Biochemistry and Physiology Part A: Physiology, 118(4), 1291–1299. 10.1016/S0300-9629(97)00243-0

Magurran, A. (2004). Measuring Biological Diversity. Wiley Blackwell Science Ltd. https://www.wiley.com/en-us/Measuring+Biological+Diversity-p-9780632056330

Morgan, K. V., Hurly, T. A., & Healy, S. D. (2014). Individual differences in decision making by foraging hummingbirds. Behavioural Processes, 109, 195–200. 10.1016/j.beproc.2014.08.015

Núñez, P., Méndez, M., & López-Rull, I. (2021). Can foraging hummingbirds use smell? A test with the amazilia hummingbird Amazila amazilia. Ardeola, 68(2), 433–444. 10.13157/arla.68.2.2021.sc2

O’Farrell, S., Sanchirico, J. N., Spiegel, O., Depalle, M., Haynie, A. C., Murawski, S. A., Perruso, L., & Strelcheck, A. (2019). Disturbance modifies payoffs in the explore-exploit trade-off. Nature Communications, 10(1), Article 1. 10.1038/s41467-019-11106-y

O’Hara, M., Mioduszewska, B., von Bayern, A., Auersperg, A., Bugnyar, T., Wilkinson, A., Huber, L., & Gajdon, G. K. (2017). The temporal dependence of exploration on neotic style in birds. Scientific Reports, 7(1), Article 1. 10.1038/s41598-017-04751-0

Oksanen, J., Simpson, G. L., Blanchet, F. G., Kindt, R., Legendre, P., Minchin, P. R., O’Hara, R. B., Solymos, P., Stevens, M. H. H., Szoecs, E., Wagner, H., Barbour, M., Bedward, M., Bolker, B., Borcard, D., Carvalho, G., Chirico, M., Caceres, M. D., Durand, S., … Weedon, J. (2022). vegan: Community ecology package (2.6-4) [Computer software]. https://cran.r-project.org/web/packages/vegan/index.html

Reader, S. M. (2015). Causes of individual differences in animal exploration and search. Topics in Cognitive Science, 7(3), 451–468. 10.1111/tops.12148

Schulz, E., & Gershman, S. J. (2019). The algorithmic architecture of exploration in the human brain. Current Opinion in Neurobiology, 55, 7–14. 10.1016/j.conb.2018.11.003

Shannon, C. E. (1948). A mathematical theory of communication. The Bell System Technical Journal, 27(3), 379–423. 10.1002/j.1538-7305.1948.tb01338.x

Shea, N., & Heyes, C. (2010). Metamemory as evidence of animal consciousness: The type that does the trick. Biology & Philosophy, 25(1), 95–110. 10.1007/s10539-009-9171-0

Sherratt, T. N., & Morand-Ferron, J. (2018). The adaptive significance of age-dependent changes in the tendency of individuals to explore. Animal Behaviour, 138, 59–67. 10.1016/j.anbehav.2018.01.025

Shettleworth, S. J. (2010). Cognition, evolution, and behavior. Oxford University Press.

Suarez, R. K. (1992). Hummingbird flight: Sustaining the highest mass-specific metabolic rates among vertebrates. Experientia, 48(6), 565–570. 10.1007/BF01920240

Tello-Ramos, M. C., Hurly, A. T., & Healy, S. D. (2019). From a sequential pattern, temporal adjustments emerge in hummingbird traplining. Integrative Zoology, 14(2), 182–192. 10.1111/1749-4877.12370

Tello-Ramos, M. C., Hurly, T. A., & Healy, S. D. (2015). Traplining in hummingbirds: Flying short-distance sequences among several locations. Behavioral Ecology, 26(3), 812–819. 10.1093/beheco/arv014

Temeles, E. J., Shaw, K. C., Kudla, A. U., & Sander, S. E. (2006). Traplining by purple-throated carib hummingbirds: Behavioral responses to competition and nectar availability. Behavioral Ecology and Sociobiology, 61(2), 163–172. 10.1007/s00265-006-0247-4

Vámos, T. I. F., Tello-Ramos, M. C., Hurly, T. A., & Healy, S. D. (2020). Numerical ordinality in a wild nectarivore. Proceedings of the Royal Society B: Biological Sciences, 287(1930), 20201269. 10.1098/rspb.2020.1269

Varma, S., Rajesh, T. P., Manoj, K., Asha, G., Jobiraj, T., & Sinu, P. A. (2020). Nectar robbers deter legitimate pollinators by mutilating flowers. Oikos, 129(6), 868–878. 10.1111/oik.06988

Ward, B. J., Day, L. B., Wilkening, S. R., Wylie, D. R., Saucier, D. M., & Iwaniuk, A. N. (2012). Hummingbirds have a greatly enlarged hippocampal formation. Biology Letters, 8(4), 657–659. 10.1098/rsbl.2011.1180

Wolf, L. L., & Hainsworth, F. R. (1990). Non-random foraging by hummingbirds: Patterns of movement between *Ipomopsis aggregata (Pursch) V. Grant* inflorescences. Functional Ecology, 4(2), 149–157. 10.2307/2389334

