## supplement for "Learning when to learn: hummingbirds adjust their exploration behaviour to match the value of information"

**Table S1. Analysis of foraging performance, or the probability that a bird probed a rewarded flower.** Estimates are reported from a binomial mixed-effects regression model (n = 3,433 probes in 34 sessions by 5 birds). This model is also shown in Figure 1D.

| Fixed effect | Estimate | SE | Test statistic | P-value |
| --- | --- | --- | --- | --- |
| Body mass (g) | 0.16 | 0.12 | 1.26 | 0.21 |
| Time (h) | 0.17 | 0.03 | 4.95 | < 0.0001 |
| Session # (1^st^ order) | –15.15 | 5.15 | –2.94 | 0.003 |
| Session # (2^nd^ order poly term) | 0.99 | 4.92 | 0.20 | 0.84 |
| Time : Session # (1^st^ order) | 8.85 | 2.02 | 4.39 | < 0.0001 |
| Time : Session # (2^nd^ order) | 4.55 | 1.98 | 2.30 | 0.02 |

**Table S2. Analysis of choice shift, or the probability that a bird changed which flower it chose from one probe to the next.** Estimates are reported from a binomial mixed-effects regression model (n = 3,399 probes in 34 sessions by 5 birds). This model is also shown in Figure 2A.

| Fixed effect | Estimate | SE | Test statistic | P-value |
| --- | --- | --- | --- | --- |
| Body mass (g) | –0.23 | 0.30 | –0.76 | 0.45 |
| Time (h) | 0.08 | 0.04 | 1.75 | 0.08 |
| Session # (1^st^ order) | 39.44 | 6.29 | 6.27 | < 0.0001 |
| Session # (2^nd^ order poly term) | –15.94 | 6.06 | –2.63 | 0.009 |
| Time : Session # (1^st^ order) | –16.99 | 2.41 | –7.05 | < 0.0001 |
| Time : Session # (2^nd^ order) | 1.19 | 2.45 | 0.49 | 0.63 |

**Table S3. Analysis of choice diversity for the most recent 5 flowers that a bird probed.** Estimates are reported from a Gaussian mixed-effects regression model (n = 3,297 probes in 34 sessions by 5 birds). This model is also shown in Figure 2B.

| Fixed effect | Estimate | SE | Test statistic | P-value |
| --- | --- | --- | --- | --- |
| Body mass (g) | –0.04 | 0.05 | –0.88 | 0.38 |
| Time (h) | 0.02 | 0.007 | 3.76 | 0.0002 |
| Session # (1^st^ order) | 10.74 | 1.07 | 10.01 | < 0.0001 |
| Session # (2^nd^ order poly term) | –5.03 | 1.00 | –5.04 | < 0.0001 |
| Time : Session # (1^st^ order) | –4.43 | 0.39 | –11.26 | < 0.0001 |
| Time : Session # (2^nd^ order) | 0.53 | 0.39 | 1.38 | 0.17 |

**Use of previously rewarded flowers**

We checked whether at the start of a new session, the birds were more likely to choose flowers they had recently experienced as rewarded during their previous (i.e., most recent) session. We focused on sessions 2-7 as there is no previous session before session 1. On their first probe of a session, the birds chose a flower that was a previously experienced reward from the most recent session 30% of the time. When examining their first five probes of a session, 38% of these were made to a previously experienced reward from the most recent session. To compare these rates to chance, we considered a “null” forager that chooses randomly within the uppermost three rows of flowers. The null expectation is previous rewards are chosen 20% of the time on average. The actual hummingbirds were significantly more likely to choose a previous reward in their first five probes as compared to this null expectation (paired t-test, p < 0.0001, n = 30 bird-sessions). When considering their first probe only, there was no significant difference between their probability of choosing a previous reward and the null expectation (paired t-test, p = 0.27, n = 30 bird-sessions). Overall, it is important to note that the relatively low frequencies observed (e.g., 38% among first five probes) demonstrates that the hummingbirds did not limit their initial visits to previously learned rewards.
